# Converting a broad matrix metalloproteinase family inhibitor into a specific inhibitor of MMP-9 and MMP-14

**DOI:** 10.1101/241372

**Authors:** Jason Shirian, Valeria Arkadash, Itay Cohen, Tamila Sapir, Evette S. Radisky, Niv Papo, Julia M. Shifman

## Abstract

MMP-14 and MMP-9 are two well established cancer targets for which no specific clinically relevant inhibitor is available. Using a powerful combination of computational design and yeast surface display technology, we engineered such an inhibitor starting from a non-specific MMP inhibitor, N-TIMP2. The engineered purified N-TIMP2 variants showed enhanced specificity towards MMP-14 and MMP-9 relative to a panel of off-target MMPs. MMP-specific N-TIMP2 sequence signatures were obtained that could be understood from the structural perspective of MMP/N-TIMP2 interactions. Our MMP-9 inhibitor exhibited 1000-fold preference for MMP-9 vs. MMP-14, which is likely to translate into significant differences under physiological conditions. Our results provide new insights regarding evolution of promiscuous proteins and optimization strategies for design of inhibitors with single-target specificities.

## Introduction

Matrix metalloproteinases (MMPs) belong to a family of over twenty MMPs and twelve homologous ADAMs proteases, enzymes that are respectively responsible for degradation of the extracellular matrix and membrane proteins [1]. MMPs differ in domain composition and size, yet all of them share a similar catalytic domain with a Zn^2+^ ion in the active site. MMPs are first produced in an inactive form and are activated by the cleavage of a pro-domain by certain MMPs or other proteases. MMP function is important in biological processes involving tissue remodeling, such as development and immune response [2].

Upregulation of MMPs or lack of MMP inhibition leads to various diseases including arthritis, chronic obstructive pulmonary disease, inflammatory bowel diseases, sepsis and various types of cancer [2]. In cancer, some MMPs play a crucial role in angiogenesis, metastasis and other aspects of tumor growth through cleavage and activation of a variety of different proteins [3–5]. The unhindered digestion of the extracellular matrix by specific MMPs such as MMP-2 and MMP-9 allows for tumor progression and cancer cells to invade and traverse from one tissue to another, resulting in the appearance of new tumors, and the activity of other MMPs, such as membrane bound MMP-14 (also called MT1-MMP) that promote cancer by activating other MMP family members [6]. Inhibition of particular MMPs could hence reverse cancer progression and reduce the spread of cancer cells.

Due to their importance to disease, MMPs have been the target for drug design efforts over the past thirty years, and several small molecules were developed early on for MMP inhibition [3]. Yet, all of them have failed in clinical trials [7]. The major reason for the failure of these drugs was their low specificity: small-molecule MMP inhibitors were designed to bind to the active-site Zn^2+^ and hence reacted with Zn^2+^ and other heavy metals in various proteins in the body and thus were highly toxic. In addition, drugs directed at multiple MMP family members elicited unexpected effects due to diverse MMP activities. In fact, some MMPs have been documented to play essential roles and even anti-tumorigenic roles [8–10], pointing to the importance of developing selective inhibitors that target only one or a narrow range of MMPs.

Such high specificity, while difficult to obtain with small molecules, could be achieved in protein-based inhibitors [11–14]. Proteins possess a greater potential for high specificity due to their large interaction surface that involves not only the highly conserved catalytic site but also more variable surrounding residues. In this respect, antibodies have been developed that specifically target MMP-9 [15] and MMP-14 [16, 17], proving the possibility of engineering a type-specific MMP inhibitor. Besides antibodies, other attractive scaffolds for MMP inhibitor design are the natural broad inhibitors of the MMP family, tissue inhibitors of metalloproteinases (TIMPs). The mammalian TIMPs include four homologous proteins (TIMP1–4) that exhibit slightly different preferences for various MMPs [18]. The advantage of using TIMPs for MMP inhibitor design is their already high affinity towards various MMPs (10^-10^-10^-9^ M), no toxicity, no immunogenicity and their smaller size that facilitates their easy production by microbial expression and supplies them with better tissue perfusion rates in comparison to antibodies. To this end, TIMPs have been already a subject of several protein engineering studies [19–22].

For this study, as a scaffold for MMP inhibitor design, we chose the N-terminal domain of TIMP-2 (N-TIMP2), consisting of 126 amino acids. The isolated N-TIMP2 remains a strong inhibitor of various MMPs [23, 24] and is more easily expressed in bacterial cultures compared to the full-length TIMP2. Most importantly, N-TIMP2 has a smaller protein interaction network compared to the full length protein; it cannot participate in interactions with the MMP hemopexin-like domain, thus losing its ability to activate certain MMPs [25]. The crystal structures of TIMP2-MMP complexes [19, 26–29] show that binding of N-TIMP2 to MMPs is mediated by four discontinuous regions (Figure 1A). These regions include the N-TIMP2 N-terminal region that comes in close proximity to the highly-conserved enzyme active site and three loops (35–42, 66–72, and 97–99). While previous TIMP protein engineering studies mostly focused on mutating the N-terminal residues, we decided to focus on the loops that interact with less conserved residues further away from the enzyme active site. Such a strategy is likely to yield inhibitors that can discriminate between different MMPs.

**Figure 1:**
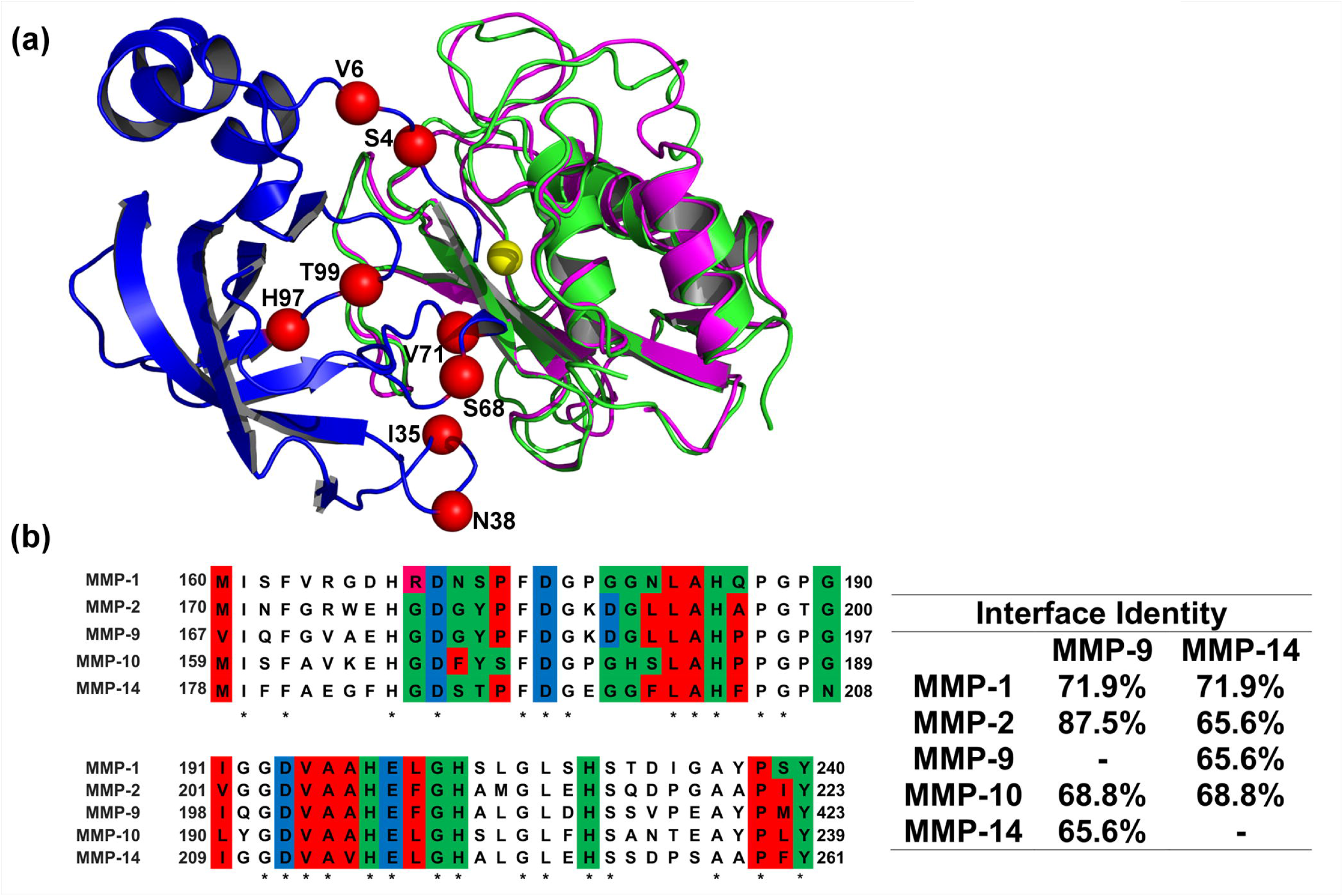
(A) Structural alignment of MMP-9_CAT_ shown in magenta (PDB: 1L6J) and MMP-14_CAT_ shown in green (PDB: 1BUV) in complex with N-TIMP2 shown in blue. Red spheres indicate positions where mutations were allowed in the designed combinatorial N-TIMP2 library and are labelled with their WT identity. Yellow sphere represents Zn^2+^ atom in the MMP active site. (B) **Left panel** A pairwise alignment of residues on MMP-1_CAT_, MMP-2_CAT_, MMP-9_CAT_, MMP-10_CAT_, and MMP-14_CAT_ (generated with the aid of EMBL Clustal Omega). Residues located in the WT N-TIMP2:MMP interface are colour coded according to chemical character of their side chains. Red represents hydrophobic residues, green represents hydrophilic residues, pink represents positively charged residues and blue represents negatively charged residues. These interface residues are at a distance of 4A or less from N-TIMP2. Asterisks indicate that there is a consensus at a particular position. **Right panel** Percent of amino-acid identity between binding interface residues for various pairs of MMPs is shown.

The goal of this work was to evolve the multispecific inhibitor N-TIMP2 into a high-affinity and high-specificity inhibitor of one out of two different MMP family members: a membrane-bound MMP, MMP-14, or the secreted gelatinase, MMP-9. The catalytic domains of MMP-14 and MMP-9 (termed MMP-14_CAT_ and MMP-9_CAT_, respectively) are structurally very similar (RMSD of 0.6 Å) showing an overall sequence identity of 45% and 66% identity among the residues that contact N-TIMP2 (Figure 1A and B). N-TIMP2 is a prototypical multispecific protein [30], whose sequence is a compromise for interactions with all MMP family members and thus is not optimal for interactions with each specific MMP [31, 32]. Recent studies on other multispecific proteins demonstrated that such proteins could be converted into highly specific binders of a particular target either through computational methods [33–35] or directed evolution approaches [36, 37].

In our previous work [38], we used the computational saturation mutagenesis protocol [35, 39] to explore the effect of all single mutations in the N-TIMP2 binding interface on N-TIMP2’s binding affinity and specificity to MMP-14_CAT_. Our computational results pointed to a high frequency of mutations that enhance N-TIMP2’s affinity and specificity toward MMP-14. Experimental results confirmed our predictions, identifying ten affinity-enhancing mutations and eleven specificity-enhancing mutations for MMP-14 relative to MMP-9. However, the enhancement in binding specificity remained relatively low for single mutations, not exceeding a factor of ten. Clearly, introduction of multiple mutations into N-TIMP2 would be required to achieve further improvements. In the present study, we thus constructed a small combinatorial library of N-TIMP2 mutants that is rich in affinity-and specificity-enhancing mutations and used yeast surface display (YSD) technology [40] to select N-TIMP2 variants that bind selectively to MMP-14_CAT_ and separately to MMP-9_CAT_. Our results show that selective N-TIMP2 inhibitors could be found and the acquired mutations contained specific signatures that define specificity for each of the MMPs.

## Materials and Methods

### Preparation of N-TIMP2 Libraries for YSD

The pCHA yeast surface display plasmid [41] was used for displaying N-TIMP2 WT and mutants on the yeast surface with a free N-terminus. The library of N-TIMP2 mutants was constructed using an assembly PCR method (see Supporting Information) randomizing positions 4, 6, 35, 38, 68, 71, 97, 99 to particular amino acids specified in Table 1. The transformation of the library to yeast was performed by simultaneously transforming the amplified mutant N-TIMP2 fragments with a pCHA[VK1] vector linearized by BamHI-HF and Nhe-HF restriction enzymes (New England Biolabs) into a competent EBY100 *Saccharomyces cerevisiae* yeast strain by electroporation.

**Table 1:** Design of the small focused N-TIMP-2 library. Desired amino acid choices display mutations that were tested experimentally and showed enhanced binding specificity of N-TIMP-2 towards MMP-14_CAT_ in our previous work when introduced as single mutations. The rightmost column shows all the amino acid choices encoded in the library at each position, including extra choices that appeared due to encoding each position with a particular degenerate codon. WT amino acid identities are underlined.

| N-TIMP-2 Position | Desired Amino Acid Choices | Degenerate Codon | Actual Encoded Amino Acids |
| --- | --- | --- | --- |
| **4** | A, Q, E, <u>S</u> | BMR | A, Q, E, P, <u>S</u>, stop |
| **6** | R, <u>V</u> | SKN | R, G, L, <u>V</u> |
| **35** | E, <u>I</u>, K | RWA | E, <u>I</u>, K, V |
| **38** | <u>N</u>, Q | MAN | <u>N</u>, Q, H, K |
| **68** | <u>S</u>, W, Y | WRS | R, N, C, K, <u>S</u>, W, Y, stop |
| **71** | N, <u>V</u> | RWY | N, D, I, <u>V</u> |
| **97** | R, <u>H</u> | CRY | R, <u>H</u> |
| **99** | <u>T</u>, Y | WMY | N, S, <u>T</u>, Y |

### FACS Analysis and Sorting

Growth of yeast (EBY100) cells, FACS analyses and sorting of libraries was performed as in Arkadash et al. [42]. For the detection of binding to MMP-14_CAT_, the cells were incubated with MMP-14_CAT_ labelled with DyLight-488 for 1 hour on ice. An identical process was performed for detection of MMP-9_CAT_ binding, where MMP-9_CAT_ is labelled with DyLight-650. Three rounds of selection were performed with lowering the concentration of the target MMP and raising the concentration of the off-target MMP (see Figure S2).

### Yeast Library Sequencing

After each round of sorting, cells collected from the flow cytometry sorter were grown in SDCAA medium (0.54% Na_2_HPO_4_, 0.86% Na_2_HPO_4_•H_2_O, 1.5% agar, 2% dextrose, 0.67% yeast nitrogen base, 0.5% Bacto casamino acids) supplemented with chloramphenicol (25 µg/ml) for ~36 hrs. 10 µl of resulting culture was diluted in SDCAA to a volume final of 2 ml, 20 µl of this diluted solution was plated on SDCAA agar plates supplemented with chloramphenicol (25 µg/ml). Single colonies were picked and served as input for colony PCR and subsequent sequencing (see Supporting Information for details).

### N-TIMP2 Expression and Purification

The N-TIMP2 mutant genes were transferred from the pCHA plasmid obtained from yeast lysates (lysed by boiling in 0.02 M NaOH) to the pPICZαA *P. pastoris* expression vector by transfer PCR [43].

PCR products were subsequently linearized with SacI (New England Biolabs) transformed into chemical competent DH5α *E. coli*. The transformed bacteria were plated on LB agar plates containing 25 µg/ml Zeocin (InvivoGen, CA, USA). After an overnight incubation at 37°C, three colonies were chosen for growth in liquid LB medium at 37°C for ~16 hours, the plasmid was purified, and the correct sequence was verified by sequencing. Bacterial cultures possessing plasmids encoding for the correct sequence were grown for ~16 hours at 37°C for large scale plasmid purification by maxiprep (Geneaid, Taiwan). The purified plasmid was then transformed to X33 *P. pastoris* yeast and recombinant protein production was induced as described in our previous work [42].

The proteins were purified as described in our previous work [42]. Protein concentrations were determined by UV-Vis absorbance at 280 nm, with an extinction coefficient (ε_280_) of 13,325 M^-1^cm^-1^ for WT N-TIMP2 and all its mutants.

### MMP Enzyme Expression and Purification

The human MMP-9 catalytic domain (MMP-9_CAT_), residues 107–215, 391–443, was expressed and purified as previously described [15], with the following modifications; the gene was expressed in Bl21(DE3)pLysS *E.coli* cells in a pET28 vector (with an N-terminal 6×His tag) and induced with 1mM isopropyl β-D-1-thiogalactopyranoside (IPTG) overnight at 30°C. The enzyme was nickel affinity purified, subsequently purified by anion exchange chromatography and then by size exclusion chromatography. The catalytic domains of MMP-1, MMP-2, MMP-10, and MMP-14 (MMP-1_CAT_, MMP-2_CAT_, MMP-1 0_CAT_ and MMP-14_CAT_, respectively) were expressed and purified as previously described [29, 42, 44].

### Enzymatic Activity Assay of N-TIMP2:MMP binding

Varying concentrations of N-TIMP2 were incubated with 0.5 nM MMP-1_CAT_, 6 nM MMP-2_CAT_, 0.1 nM MMP-9_CAT_, 0.5 nM MMP-10_CAT_, 0.2 nM MMP-14_CAT_ under conditions similar to that described in [42]. Fluorescence resulting from cleavage of fluorogenic substrate was measured at 395 nm, at least every ten seconds for at least 6 minutes, immediately after addition of fluorogenic substrate with irradiation at 325 nm on a Biotek Synergy H1 plate reader (BioTek, VT, USA). At least three assays were performed for each N-TIMP2 protein.

### 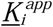 Determination

Initial velocities of the enzymatic (MMP) cleavage of the fluorogenic substrate were derived from the fluorescence generated by this cleavage. Fraction of MMP activity was determined from the enzymatic assay, expressed as the inhibited velocity (V_i_) divided by the uninhibited velocity (V_0_), and was plotted against the corresponding concentration of N-TIMP2. From this plot the apparent 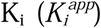 value was solved for based on the following equation [45]:

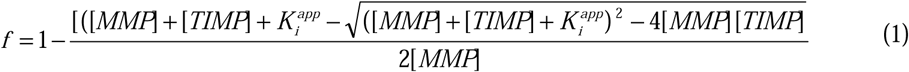

*f=* fraction of activity, V_i_/V_0_

Specificity for MMP-X relative to MMP-Y was calculated according to the following formula:

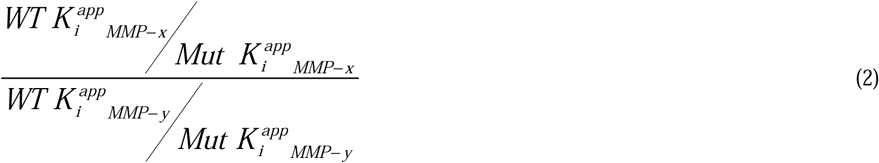

### Structural Modeling

Modeling of N-TIMP2 in complex with MMP-14_CAT_ and MMP-9_CAT_ was performed as in our previous study [38]. For modeling the mutations in the N-TIMP2 binding interface, the energy function optimized by Sharabi et al. [38, 46] was used. All mutations present in each N-TIMP2 mutant were simultaneously introduced into N-TIMP2 first with no backbone flexibility and then using backbone flexibility using the RosettaBackrub server [47]. The output of this protocol was a multiple-mutant structure along with its △△G_bind_ value. The difference in molecular interactions due to introduction of each mutation was analyzed in the context of both structures, the N-TIMP2/MMP-14_CAT_ and the N-TIMP2/MMP-9_CAT_ structures to determine which type of interactions cause a specificity switch.

## Results

N-TIMP2 library construction To obtain inhibitors that bind to either MMP-14_CAT_ or MMP-9_CAT_ with high affinity and high specificity, we constructed a small computationally designed combinatorial library that mutated only a few positions on the N-TIMP2 binding interface. We randomized eight positions in the direct N-TIMP2 binding interface (Figure 1A) to a limited set of amino acids, including either WT choice or mutation(s) observed to enhance binding affinity and/or specificity towards MMP-14_CAT_ in our previous study [38] (Table 1). Additional amino acid choices were included in the library when no degenerate codon could be found to simultaneously encode for only the desired choices. The designed library included 68480 different full-length N-TIMP2 sequences, and was enriched in sequences with enhanced specificity towards MMP-14_CAT_. Sequencing of the library before selective sorting revealed that the clones in the library contained on average of 6±2 mutations per gene.

**Selection of high specificity MMP-9 and MMP-14 binders**. YSD technology was used to select specific binders for MMP-9_CAT_ and MMP-14_CAT_ in two parallel experiments. The N-TIMP2 library was expressed on the surface of yeast cells using the pCHA expression vector, where the yeast protein Aga2p is fused to the C-terminus of N-TIMP2, leaving the N-terminus of the N-TIMP2 free for interaction with the target MMPs (Figure 2A). This arrangement is crucial since the N-terminus of N-TIMP2 mediates binding to the MMP active site and thereby facilitates proper inhibition of MMPs, as could be inferred from the crystal structure of N-TIMP2/MMP complexes and was confirmed in previous experiments [42, 48–50]. The c-Myc tag was used to monitor protein expression, through binding of an anti-c-Myc antibody and a fluorescently labelled secondary antibody against the anti-c-Myc. In our selection experiments, we used both positive and negative selection, by simultaneously incubating the N-TIMP2 libraries with two targets, MMP-14_CAT_ and MMP-9_CAT_, labelled with two different fluorescent dyes, Dylight-488 and Dylight-650, respectively (Figure 2A). All sorts were performed using fluorescence activated cell sorting (FACS) (Figure 2B, C). To eliminate N-TIMP2 mutants that are not expressed (either due to deletions or due to frameshifts in the N-TIMP2 gene) we performed an initial pre-sorting round, by selecting the cells above a certain expression signal (Figure 2B).

**Figure 2:**
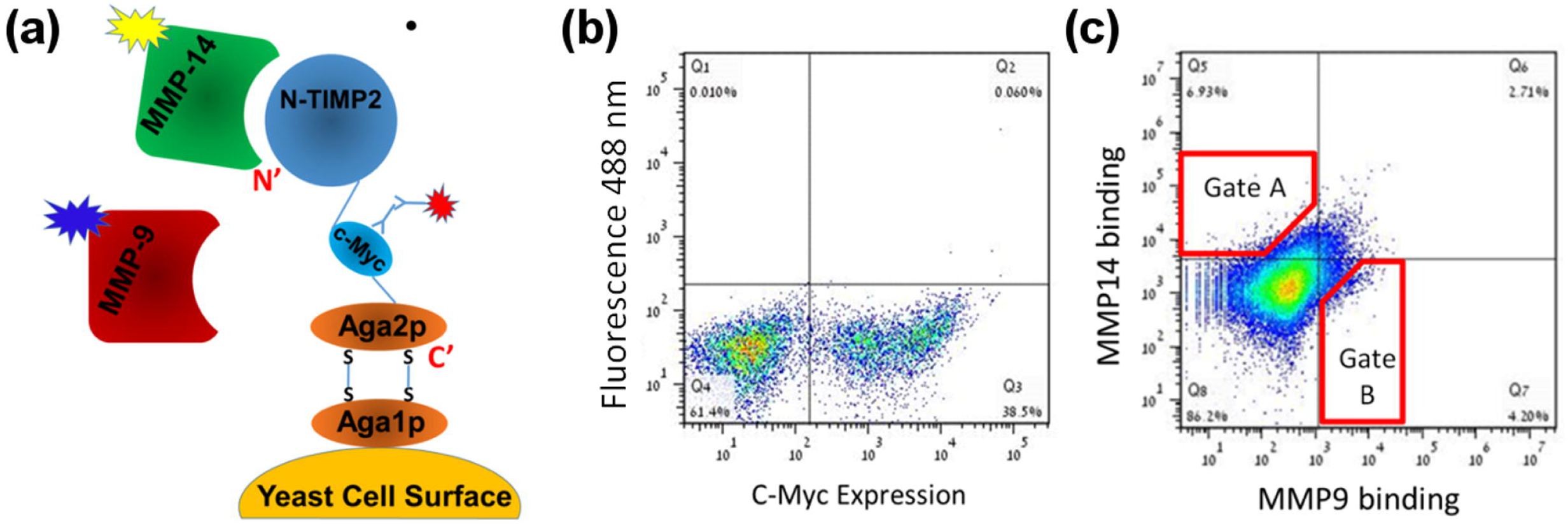
(A) Schematic representation of N-TIMP2 displayed on the yeast surface (adapted from Chao et al.[56]). The N-terminus of N-TIMP2 is free and facing away from the yeast surface. (B) Expression of the construct was monitored by FACS by fluorescently labelled antibodies (red burst in panel A) that bind the c-Myc tag in the construct. (C) Binding of N-TIMP2 to catalytic domains of MMP proteins in solution was monitored by FACS by incubating the N-TIMP2 library with two MMP targets (MMP-9_CAT_) and (MMP-14_CAT_) simultaneously and using two different fluorescent labels for each MMP proteins (yellow and blue bursts in panel A). Red polygons represent sample cell collection gates, A for MMP-14_CAT_ and B for MMP-9_CAT_. The figure here was produced in the initial sorting round on the focused library.

Following this initial sorting, the libraries were incubated with 500 nM of MMP-14_CAT_ and 50 nM of MMP-9_CAT_ (Figure 2C). A lower concentration of MMP-9_CAT_ compared to MMP-14_CAT_ was necessary since N-TIMP2 has a higher natural affinity for MMP-9 as opposed to MMP-14 [38]. Under these conditions a distributed binding pattern was observed, where cells bound only MMP-14_CAT_, only MMP-9_CAT_ or both MMP-14_CAT_ and MMP-9_CAT_ (Figure 2C). We further performed three rounds of sorting each time collecting 1% of all cells exhibiting the highest binding to MMP-14 (Supplementary Figure S1, upper panel) and separately to MMP-9 (Supplementary Figure S1, lower panel). From one round to the following, the concentration of the desired MMP was reduced while the concentration of the undesired MMP was raised, to make the binding specificity selection more stringent. After each round of sorting, ~20 colonies from both selections were randomly chosen for sequencing. Sequencing results showed an increasing convergence of certain mutations at particular N-TIMP2 positions with each round of selection. After the final round of selection, we obtained the specificity signatures for N-TIMP2 binding to MMP-14_CAT_ and to MMP-9_CAT_ (Figure 3A, B). The most outstanding result for the MMP-14_CAT_ selection is a unanimous consensus to asparagine at position 71 that replaces a WT valine (Figure 3A). The sequences of N-TIMP2 mutants in the MMP-9_CAT_ selection show a substantial consensus at position 4 for proline as opposed to the WT serine as well as a mutation from valine to isoleucine at position 71. At position 38, glutamine predominates the MMP-14_CAT_ selection, but a wild-type asparagine appears most frequently in the MMP-9_CAT_ selection. Positions 4, 38, and 71 hence, stand out as the most important specificity determining positions for MMP-14 and MMP-9 binding.

**Figure 3:**
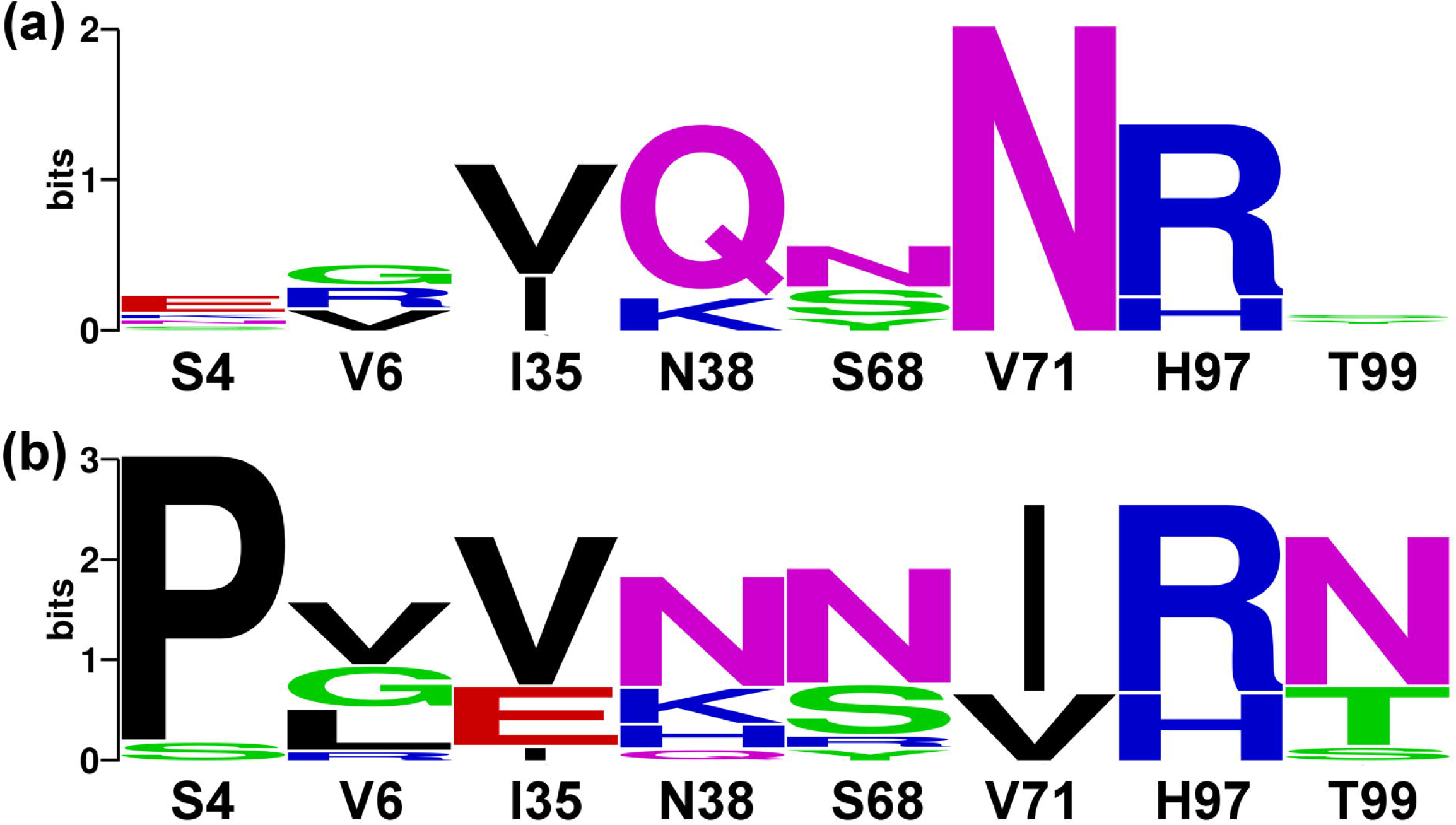
Logos based on sequencing of evolved N-TIMP2 libraries after 3 rounds of binding selection for MMP-14_CAT_ (A) and MMP-9_CAT_ (B). The logos were made through http://weblogo.berkeley.edu/logo.cgi. The WT identity of each position is shown below each logo and the size of the amino acid corresponds to its frequency among the selected sequences. The total height of the stack represents conservation at that position. Green, purple, blue, red and black letters respectively represent polar, neutral, basic, acidic and hydrophobic amino acids. Only eight binding interface positions on N-TIMP2 are shown. Additional unintended mutations were observed in more than 50% of the sequences, at an average frequency of one and a half mutations per sequence.

**Understanding specificity signatures through modeling** To understand the nature of enhanced specificity, we performed structural modeling of the engineered N-TIMP2 mutants when interacting with MMP-14 and MMP-9. At position 4, we see a strong preference for glutamate in the MMP-14_CAT_ selection while proline clearly predominates the MMP-9_CAT_ selection. Our modeling shows that a glutamate at position 4 can form an intermolecular hydrogen bond with N231 on MMP-14_CAT_ but not with MMP-9_CAT_ (Figure 4A). A proline at position 4 fits better in the more hydrophobic environment of MMP-9_CAT_ but causes a steric clash with MMP-14_CAT_, which could be removed only through substantial backbone conformational changes (Figure 4A). At position 38, glutamine predominates in the MMP-14_CAT_ selection, but a wild-type asparagine appears most frequently in the MMP-9_CAT_ selection. According to our modeling, Q38 makes a favorable hydrogen bond with N208 on MMP-14_CAT_, but cannot make such a bond with the corresponding G197 on MMP-9_CAT_ (Figure 4B). It should be noted that glycine is conserved at this position in all the MMPs tested in this work except for MMP-14_CAT_ where this position is occupied by asparagine (Figure 1B). In the MMP-9_CAT_ selection, mutation to isoleucine predominates at position 71. This isoleucine packs against hydrophobic residues on MMP-9_CAT_, namely F192 and Y179. Position 71 is highly enriched for a mutation to asparagine in the MMP-14_CAT_ selection; the asparagine could make a favorable hydrogen bond with Y203 on MMP-14_CAT_ (Figure 4C).

**Figure 4:**
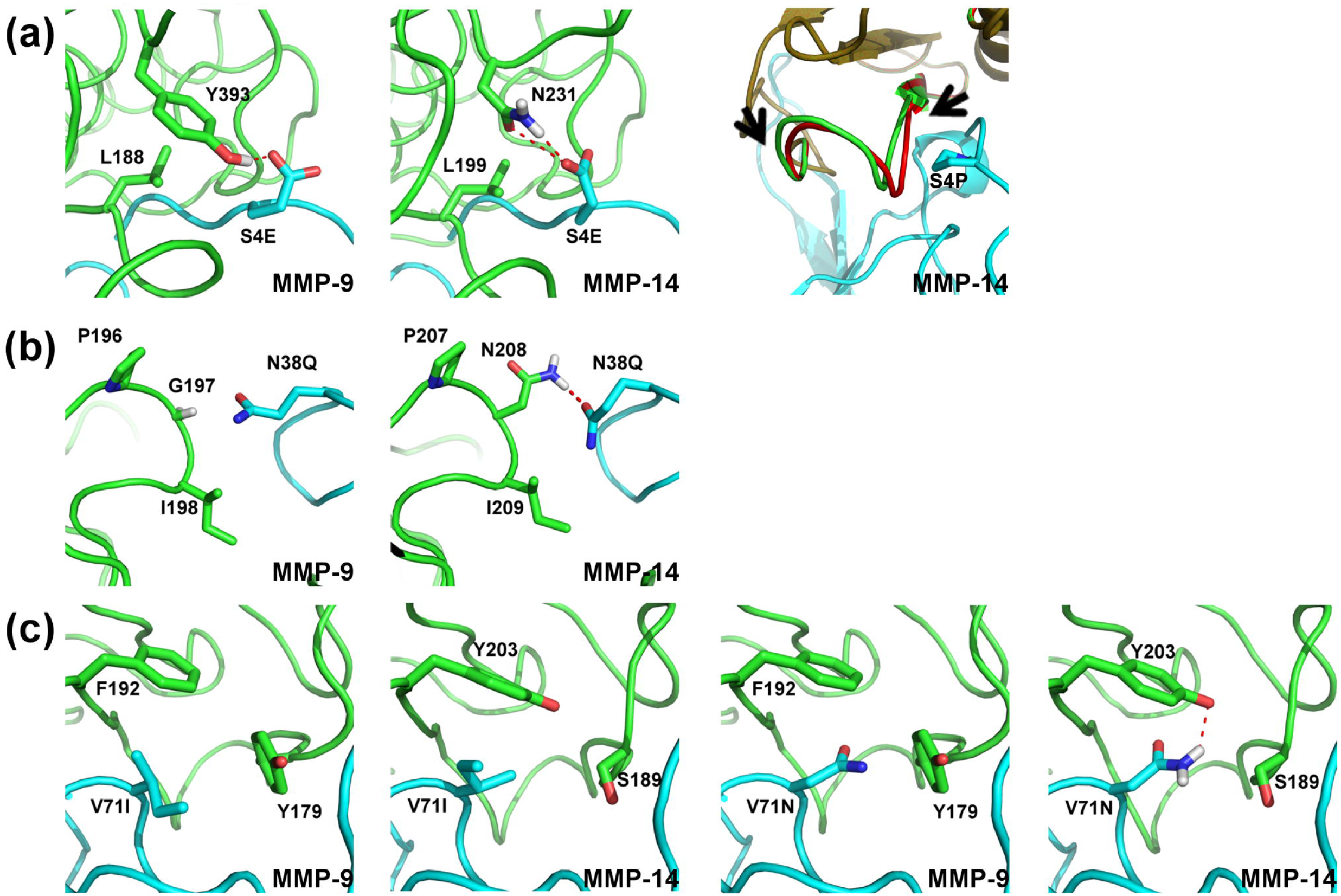
Interactions observed in computationally modeled structures of specificity enhancing mutations of N-TIMP2. N-TIMP2 is depicted in cyan and MMP-9_CAT_ or MMP-14_CAT_ is depicted in green; polar contacts are shown as red dotted lines. MMP residues shown in sticks are not further than 4Å away from the highlighted N-TIMP2 residue. Hydrogen atoms are shown as necessary for depiction of hydrogen bonds. (A) The glutamate at position 4 interacting with MMP-9_CAT_ (leftmost panel) and MMP-14_CAT_ (middle panel). Rightmost panel depicts backbone movement necessary in MMP-14_CAT_ to accommodate for proline occupying position 4 on N-TIMP2 as seen from structural alignment before and after modeling with backbone flexibility. The MMP-14_CAT_ structure before introduction of backbone flexibility is shown in red and after is shown in green, N-TIMP2 is shown in cyan. Arrows indicate region within which backbone was altered. (B) Interactions of Gln at position 38 with both MMPs are shown (C) Interactions of Ile and Asn at position 71 in the context of both MMPs are shown.

**Purification and characterization of MMP-9**_CAT_ **and MMP-14**_CAT_ **inhibitors**. To verify the success in binding specificity enhancement, six random N-TIMP2 mutants from each of the selection experiments were individually assessed for binding to MMP-14_CAT_ and MMP-9_CAT_ while displayed on the yeast surface (Figure 5A and C). The specificity of each mutant was evaluated by determining the ratio of the binding signal for the target MMP to that of the off-target MMP and then dividing by the same ratio for WT N-TIMP2. All tested mutants showed higher specificity towards the target MMP compared to that of WT N-TIMP2. In the MMP-14 selection experiment, we selected mutants 14D-4 and 14D-6 for further purification and characterization since they exhibited the highest specificity signal (Figure 5A). In the MMP-9 selection experiment, we chose mutants 9D-2 and 9D-6 since mutant 9D-2 exhibited the highest specificity signal (Figure 5C), whereas mutant 9D-6 recurred 3 out of 15 times among the sequenced clones. Table 2 lists the mutations present in the N-TIMP2 mutants chosen for purification and characterization from the designed library.

**Table 2:** Sequences of the N-TIMP2 mutants chosen for expression and purification. Mutations in N-TIMP2 variants chosen to be assayed *in vitro* as purified proteins from the small designed combinatorial focused N-TIMP2 library. “--” indicates that the WT identity at this particular position was maintained. Mutant 14D-6 contains a mutation, V18M, which was not in initial design of the library.

| Position: | **4** | **6** | **35** | **38** | **68** | **71** | **97** | **99** | Additional mutations |
| --- | --- | --- | --- | --- | --- | --- | --- | --- | --- |
| **WT** | **S** | **V** | **I** | **N** | **S** | **V** | **H** | **T** |  |
| **14D-4** | **E** | **--** | **--** | **Q** | **--** | **N** | **R** | **S** |  |
| **14D-6** | **E** | **R** | **V** | **Q** | **N** | **N** | **R** | **--** | **V18M** |
| **9D-2** | **P** | **--** | **V** | **--** | **N** | **I** | **R** | **--** |  |
| **9D-6** | **P** | **G** | **V** | **--** | **N** | **I** | **--** | **N** |  |

The four selected N-TIMP2 mutants and the WT N-TIMP2 were expressed and purified from *Pichia pastoris* yeast cells, which secrete the properly folded N-TIMP2 protein (Supplementary Figure S2)[50, 51]. The 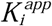 of the inhibition of MMP-14_CAT_ and MMP-9_CAT_ by N-TIMP2 mutants was determined using an MMP enzyme activity assay in the absence and the presence of the N-TIMP2 inhibitor as previously described (Figure 5B and D) [38]. Table 3 summarizes the 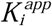 values obtained for each N-TIMP2 mutant, for inhibition of MMP-14_CAT_ and MMP-9_CAT_. All tested mutants showed a substantial increase in binding specificity towards the target MMP (Table 3 and 4). The best mutant from the MMP-14_CAT_ selection, 14D-6, with 8 mutations exhibited a specificity shift of ~60-fold. This specificity increase is due to ~3-fold improvement in affinity to the target MMP-14 and ~20-fold decrease in affinity to MMP-9 relative to WT N-TIMP2. The two expressed N-TIMP2 mutants from the MMP-9_CAT_ selection show similar double digit specificity enhancements values. The best mutant, 9D-6, exhibits a 44-fold increase in binding specificity to MMP-9_CAT_ over MMP-14_CAT_, but a slightly worse affinity to MMP-9 compared to WT N-TIMP2. Thus, the specificity increase in this mutant comes through a great reduction in affinity to MMP-14_CAT_ relative to WT N-TIMP2. Note that the 9D-6 mutant exhibits nearly 1000-fold better binding affinity towards MMP-9_CAT_ vs. MMP-14_CAT_ (Table 3). Such high affinity differences, should be significant for preferential inhibition of MMP-9 but not MMP-14 in cellular environments.

**Table 3:** Inhibitory activities of the purified proteins. Purified N-TIMP2 mutants were assayed for their ability to inhibit MMP-14_CAT_ and MMP-9_CAT_. 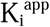 values were obtained from at least three such assay experiments for each protein and averaged, each value is reported as 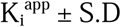. The exceptions are a few experiments where very high mutant concentration was required and thus only one experiment was performed. The boldface values of each cell shows the relative inhibitory affinity for the target vs. offtarget MMP: 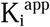 for /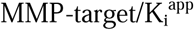 for MMP-x, where target is MMP-14_CAT_ or MMP-9_CAT_ and x is any off-target MMP.

| **$K_i^{app}$ of binding (nM)** |  |  |  |  |  |
| --- | --- | --- | --- | --- | --- |
|  | **MMP-14<sub>CAT</sub>** | **MMP-9<sub>CAT</sub>** | **MMP-1<sub>CAT</sub>** | **MMP-2<sub>CAT</sub>** | **MMP-10<sub>CAT</sub>** |
| **WT** | 2.52±0.32 | 0.12±0.02 | 0.25±0.02 | 2.65±0.76 | 4.3±0.29 |
| **14D-4** | 1.02±0.62 | 1.28±0.18 | 0.31±0.03 | 83.96±4.57 | 78.91±41.55 |
|  | **1.0** | **0.80** | **3.38** | **0.01** | **0.01** |
| **14D-6** | 0.89±0.06 | 2.43±0.90 | 0.78±0.21 | 224.5 | 446.35±167.09 |
|  | **1.0** | **0.37** | **1.14** | **0.004** | **0.002** |
| **9D-2** | 105.73±4.78 | 0.15±0.04 | 0.11±0.02 | 3.53±0.34 | >300 |
|  | **0.001** | **1.0** | **1.36** | **0.04** | **<0.0005** |
| **9D-6** | 463.50±33.16 | 0.50±0.12 | 9.53±3.21 | 38.89±2.50 | >300 |
|  | **0.001** | **1.0** | **0.05** | **0.01** | **<0.002** |

**Table 4:**
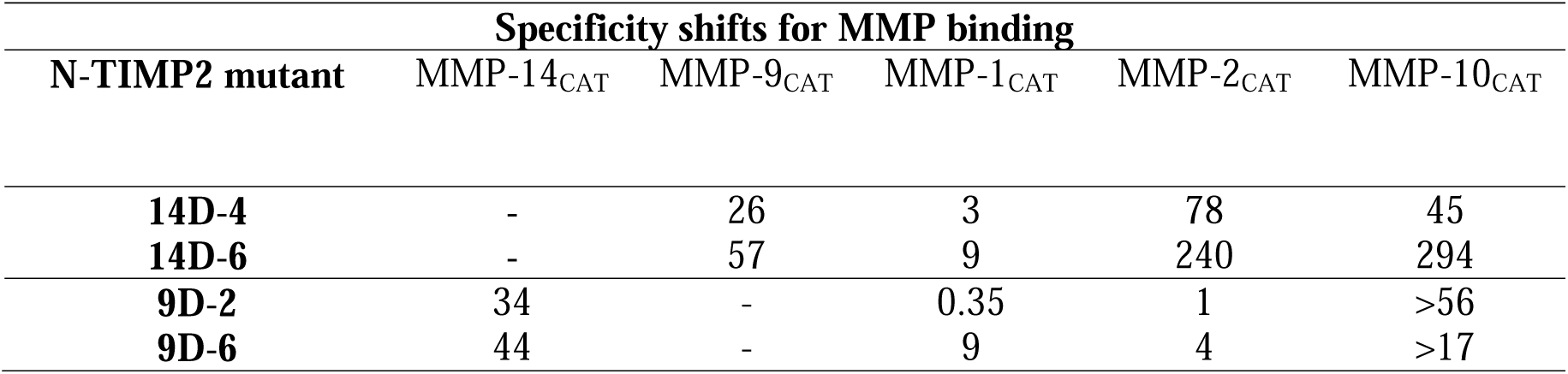
Specificity shifts of N-TIMP2 mutants in favor of MMP-14_CAT_ or MMP-9_CAT_ as compared to off-target MMPs. Specificity shifts were calculated according to eq. 2. The specificity shifts were calculated relative to MMP-14_CAT_ for the mutants engineered to bind to MMP-14_CAT_ and were calculated relative to MMP-9_CAT_ for the MMP-9_CAT_ engineered mutants.

**Figure 5:**
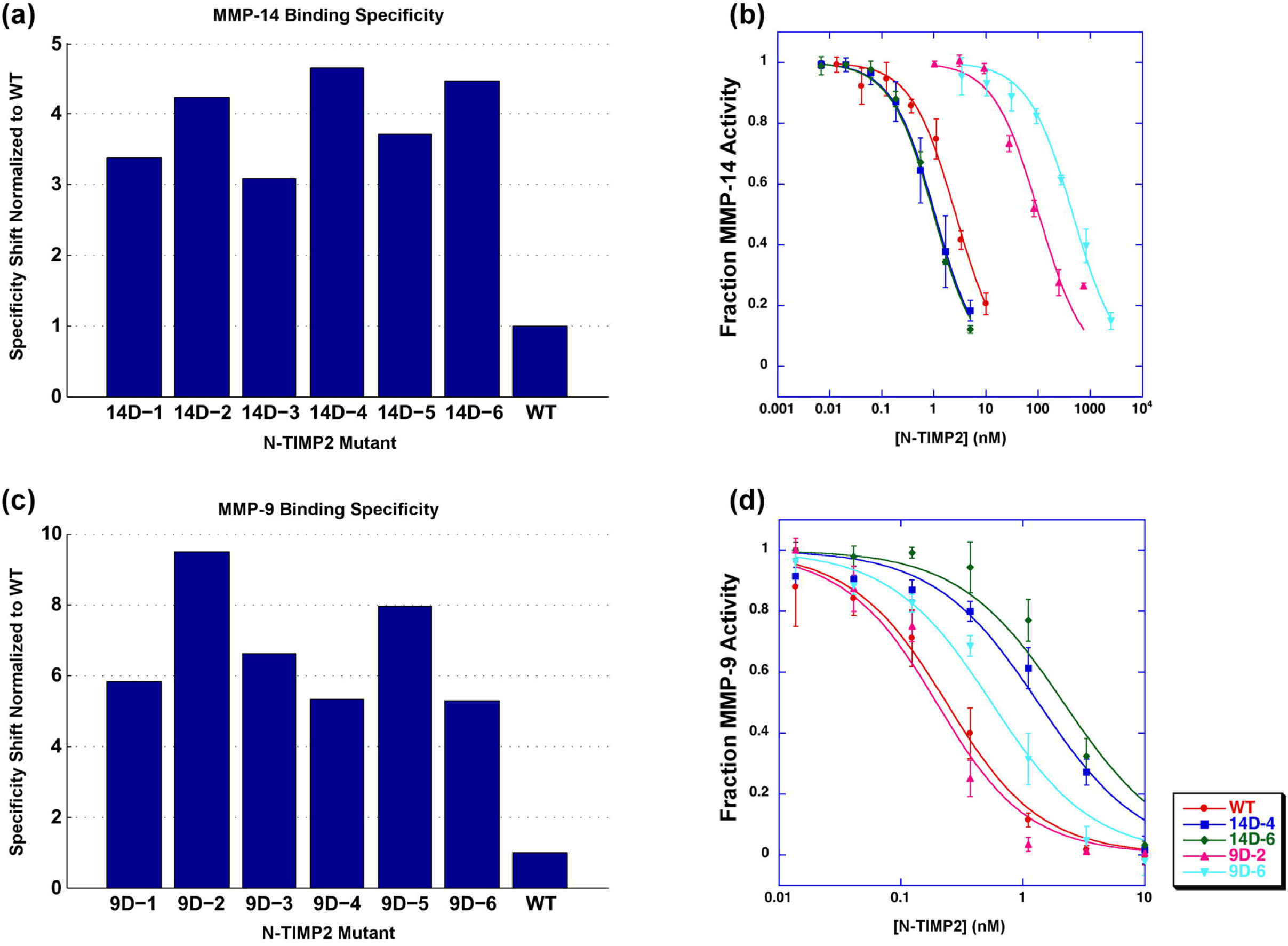
(A, C) Binding specificity of selected N-TIMP2 mutant yeast colonies assessed by FACS. Binding is determined by the average signal of fluorescence of yeast cells caused by fluorescently labelled MMP. Binding specificity is determined in the following manner in the case of MMP-14_CAT_: (MMP-14_mut_/MMP-14_wt_)/(MMP-9_mut_/MMP-9_wt_), and vice-versa for MMP-9_CAT_. (B, D) Enzyme activity assays with inhibition by WT N-TIMP2 or mutants expressed in *P. pastoris*. Data points represent an average of at least three experiments. Curves were fit to *equation 1*. (B) Enzyme activity experiments performed with MMP-14_CAT_. (D) Enzyme activity experiments performed with MMP-9_CAT_.

To determine general binding specificity of our engineered MMP-14_CAT_ and MMP-9_CAT_ inhibitors, we measured their ability to inhibit a panel of additional MMP types, MMP-1_CAT_ (collagenase), MMP-2_CAT_ (gelatinase) and MMP-10_CAT_ (stromelysin). Table 4 shows that engineered N-TIMP2 mutants generally show reduced affinity to off-target MMPs that have not been used in our selection experiments. Higher specificity shifts are observed in the MMP-14_CAT_ selection, where mutant 14D-6 exhibits a 294-fold preference for MMP-14_CAT_ over MMP-10_CAT_. The N-TIMP2 mutants from the MMP-9_CAT_ selection, especially 9D-6, are superior in absolute values of 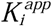, able to inhibit MMP-9_CAT_ at least 100 times better relative to all measured off-target MMPs (Table 3).

## Discussion

Designing inhibitors that act against a single or narrow range of MMPs is clearly important for targeting MMPs in diseases and much effort in this direction has been recently reported [52]. In this work, we attempted to engineer such an inhibitor starting from the natural broad MMP inhibitor, N-TIMP2. Previously, we showed that the N-TIMP2/MMP binding interface is not optimized and contains many affinity- and specificity-enhancing mutations [38]. Specifically, positions 4, 35, 38 and 68 on TIMP were determined to be cold spots of binding [53], i.e. positions where several different mutations lead to increase in binding affinity towards MMP-14. We also determined that single mutations on N-TIMP2 in general, do not increase binding specificity by more than one order of magnitude. Here, we demonstrate that incorporation of 5–8 mutations into the N-TIMP2 sequence results in two orders of magnitude enhancement in binding specificity towards the target MMP. Among the combination of mutations found in this study, we frequently observe single mutations, previously identified as specificity-enhancing (e. g. at positions 4, 38, and 71). However, beneficial single mutations, when incorporated together, result in negative cooperativity, producing smaller specificity shifts for multiple mutants compared to the sum of shifts for single mutations.

As in many other engineering studies, the N-TIMP2 mutants reported here showed reduction of binding not only to the MMP used as a competitor but also to off-target MMPs not used in the selection procedure: MMP-1_CAT_, MMP-2_CAT_, and MMP-10_CAT_. The amount of affinity reduction to off-target MMPs correlates well with the interface sequence identity between the target and the off-target MMP (Figure 1B), exhibiting the lowest specificity shift against MMP-1_CAT_ in the MMP-14_CAT_ selection experiment and MMP-1_CAT_ and MMP-2_CAT_ in the MMP-9_CAT_ selection experiment. These results demonstrate that when similarity of off-targets is highest, the specificity for the target MMP is more difficult to achieve. Furthermore, the relatively small library of N-TIMP2 mutants, containing only ~70,000 mutants, could be used to evolve this protein to interact specifically with both MMP-14 and MMP-9. The same library is likely to produce specific binders for other MMPs, opening the way to future engineering studies.

The reported N-TIMP2 mutants exhibit specificity shifts similar to what has been observed in previous N-TIMP2 engineering studies. In our recent study, N-TIMP2 was engineered using a much larger library of 10^8^ N-TIMP2 mutants and a more stringent selection procedure that included only the desired target, MMP-14_CAT_ [42]. In that work, 880-fold improvement in binding affinity for MMP-14_CAT_ was obtained, but small improvements in affinity were also reported for all but one of the off-target MMPs (MMP-1_CAT_, MMP-2_CAT_, and MMP-8_CAT_ but not MMP-10_CAT_). In another study by Brew and colleagues, N-TIMP2 was engineered to bind with high specificity to MMP-1_CAT_, and demonstrated 7fold weaker affinity to MMP-1_CAT_, no detectible binding to MMP-3_CAT_ and MMP-14_CAT_ and intermediate affinities to several other MMPs [22]. Together, these studies prove that N-TIMPs could be fine-tuned for better interaction with any MMP type, yet simultaneously discriminating against all off-target MMPs might be difficult due to relatively high sequence homology (65–80%) of various MMPs in the TIMP binding interface region. However, such discrimination might not be necessary for the development of anti-MMP drugs since the MMP inhibition pattern in real physiological environments depends on relative concentrations and localization of each MMP. Thus, each engineered N-TIMP2, with enhanced specificity and a certain set of binding affinities for each off-target MMP, might prove useful under particular MMP expression patterns.

We report for the first time, N-TIMP2 mutants engineered for MMP-9_CAT_ binding that exhibits 1000-fold better 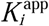 for MMP-9_CAT_ vs. MMP-14_CAT_ and large discriminations against other tested MMPs. Such a mutant is likely to selectively inhibit MMP-9 under physiological conditions and thus could serve to probe the role of MMP-9 in various diseases. As several of our previously reported MMP-14-specific N-TIMP2 variants have already showed high potency in inhibiting cancer-related MMP activity in various cellular assays [42], the MMP-14- and MMP-9-specific mutants reported in this study present interesting candidates for further testing in cellular and *in vivo* studies. Overall, our results help to understand the molecular origins of binding specificity in TIMP/MMP interactions and facilitate development of cancer drugs and diagnostic tools targeting MMPs.

## Acknowledgements

We thank M. Lebendiker for help with protein purification and A. Zika for help with FACS experiments.

